# Chromatin domains are encoded and molded by isochores: Isochores encode chromatin domains and chromosome compartments

**DOI:** 10.1101/096487

**Authors:** Giorgio Bernardi

## Abstract

The formation of mammalian chromatin domains was investigated by analyzing the domain/isochore connection. This showed that LADs correspond to GC-poor isochores and are compositionally flat, flexible chromatin structures because of the local nucleosome depletions associated with the presence of oligo-A’s. In contrast, TADs correspond to GC-rich isochores that consist of single or (much more frequently) multiple, GC peaks that shape the single or multiple, loops of TADs. Indeed, the increasing nucleosome depletions linked to the GC gradients of isochore peaks lead to an increasing chromatin flexibility (accompanied by an increasing accessibility and decreasing supercoiling). In conclusion, isochores not only encode but also mold chromatin architecture; while architectural proteins play a role in closing and insulating TAD loops. An extension of this model concerns the encoding of open and closed chromosome compartments by alternating GC-rich and GC-poor isochores, the interactions among compartments defining the 3-D chromosome folding.

---

In interphase nuclei, chromatin comprises two sets of domains that are largely conserved in mammals: LADs, the lamina associated domains (~0.5Mb median size), that correspond to GC-poor sequences (1), and TADs, the topologically associating domains (0.2-2 Mb in size), a system of GC-rich loops (2); many TADs can be resolved into contact domains (0.185 Mb median size; 3).

In spite of the recent, impressive advances in our understanding of chromatin structure, the problem of the formation mechanism(s) of LADs and TADs is still unsolved. Three interesting models proposed for TAD formation are: the “handcuff model” (4); the “extrusion model” (5,6); and the “insulation-attraction model” (2). In this article the problems of these models will be discussed and a new model will be presented.

The issue of chromatin domain formation was approached here by taking into account the observation that isochores make up the framework of chromatin domains (7) and by having a new look at isochore structure and at the isochore/chromatin domain connection. The isochore connection was also used to investigate another open problem, that of “spatial compartments” (8). This approach was mainly applied to human chromosome 21 which was chosen because this chromosome is a good representative of human chromosomes, allowing an extension of the results obtained to all other chromosomes that had been investigated previously (7).

It should be recalled that isochores, the long (>200 Kb), compositionally “fairly homogeneous” DNA sequences that belong to a small number of families (L1, L2, H1, H2, H3 in the human genome) characterized by increasing GC levels, were discovered (9) using compositional genomics, namely DNA fractionation according to the frequencies of short sequences (10; or GC levels as a proxy), the sequences that determine the fine structure of the DNA double helix and the interactions of DNA with proteins. The aim of this strategy was to understand the large-scale organization of the vertebrate genome (see ref. 11).

## The isochore/chromatin connection

The compositional profiles of the DNA sequence from human chromosome 21 were looked at through non-overlapping 100Kb windows (Fig.1A), and by using a fixed-window (Fig.1B), or a sliding window approach (Fig.1C). In Fig.1B, the compositional fluctuations of Fig.1A disappear because of the strict averaging procedure applied, whereas in Fig.1C the fluctuations are flattened but can still be seen as spikes in GC-rich regions.

**Figure 1.**
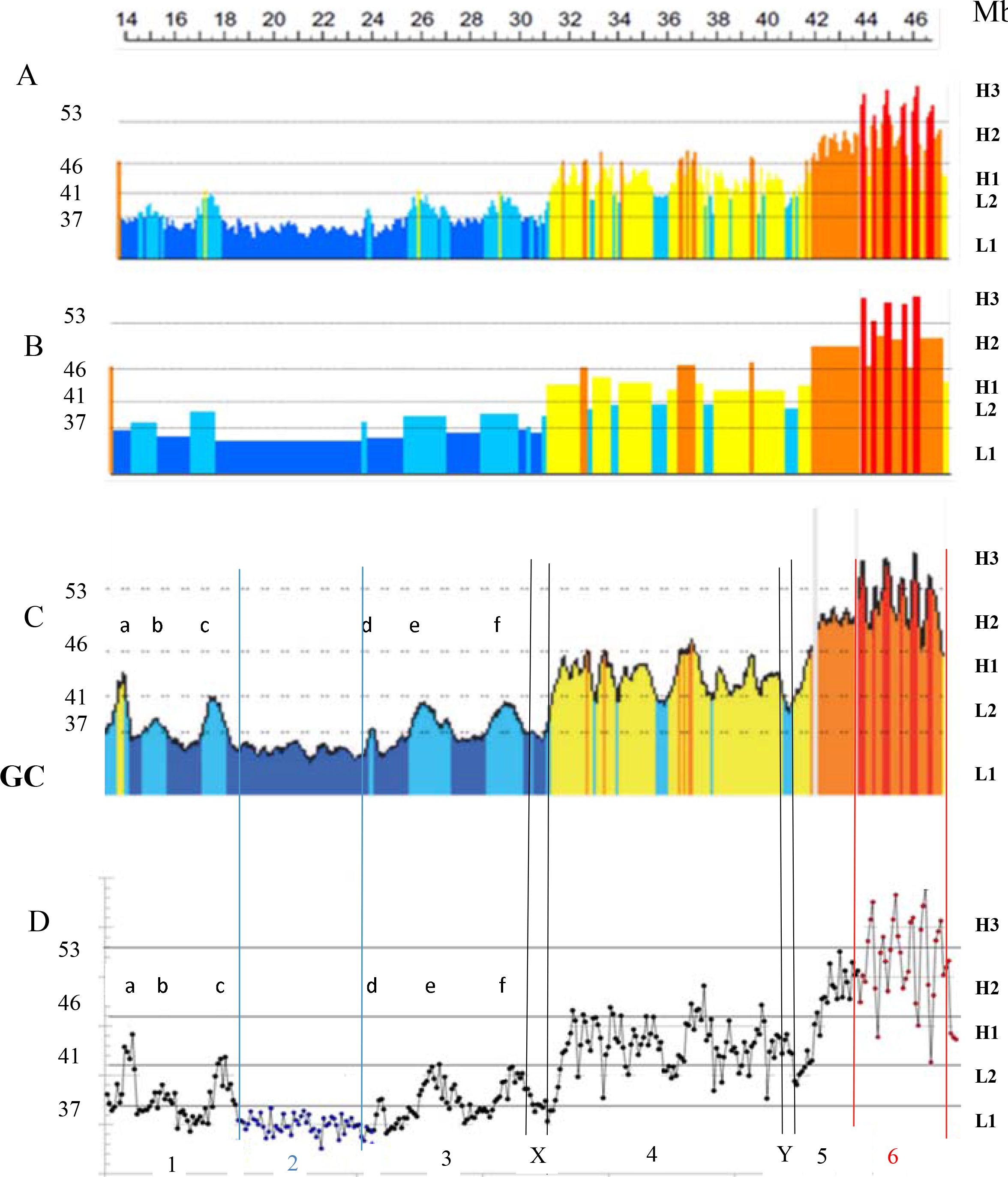
**A:** Compositional profile (38) of human chromosome 21 (release hg38) as seen through non-overlapping 100-Kb windows. DNA sequences from isochore families L1 to H3 are represented in different colors, deep blue, light blue, yellow, orange, red, respectively. The left-side ordinate values are the minima GC values between the isochore families (see Supplementary Table S1). **B.** Isochore profile of human chromosome 21 (release hg38) using a non-overlapping 100Kb window and the isoPlotter program (38).**C**. Isochore profile of human chromosome 21 using the matched b37 assembly of ref. 3 and a sliding window approach (39) with “fixed” isochore borders (see Supplementary Table 1). This profile is slightly more extended on the centromeric (left) side than those of Figs. 1A and 1B.**D**. GC levels of 100Kb windows of human chromosome 21; data of Fig. 1C are plotted as points (see Supplementary Fig.S1 for an enlarged version). This figure shows that individual isochores from the L2^+^ to H3 families are in the form of peaks that do not appear in a clear way in the standard presentation of compositional profiles of chromosomes (Fig. 1A), except for the broad, isolated H1 (a) and L2^+^ peaks (b to f). Black, blue and red lines, as well as double lines (X,Y), separate regions 1 to 6.

A new, simpler, in fact elementary, approach was used here, namely plotting the GC values of 100Kb DNA blocks as individual points (see Fig. 1D). This approach (which is preferable to that of Fig.1A for reasons of graphical clarity) was suggested by the fact that the frameworks of LADs and TADs are made up by GC-poor and GC-rich isochores, respectively (7). One may therefore imagine a possible topological similarity between the flat structure of LADs and the loops of TADs on the one hand, and the compositionally flat GC-poor and the compositionally heterogeneous GC-rich isochores on the other.

Fig. 1D (and its enlarged version, Supplementary Fig. S1) expectedly showed a compositionally flat L1 region 2 and the H1 and L2 peaks (a to f) of regions 1 and 3, that were already evident in Figs. 1A, 1B and 1C. It also led, however, to the discovery that the sequences of isochores from H1 (region 4), H2 (region 5) and H3 (region 6) families were not randomly fluctuating within the compositional borders of the corresponding families (see Supplementary Table S1), but consisted, in fact, of sets of GC peaks. Upon close inspection, these very evident peaks were seen to correspond to the minute peaks of Fig. 1C.

## Isochores and LADs

As shown in Fig. 2A, the major LAD of chromosome 21 corresponds to the large L1 isochore of region 2. In L1 isochores (~65% AT) trinucleotides only consisting of A’s and T’s represent 30% of the sequence (12) and hexa-, hepta- and octa–A’s are present (V.Sabbia, personal comm.); this situation favors local nucleosome depletions (13). The other LADs correspond to the L1 isochores that separate two L2 peaks, and to a “valley” L2 isochores comprised between two H1 isochores (on the right side of Fig. 2A).

**Figure 2.**
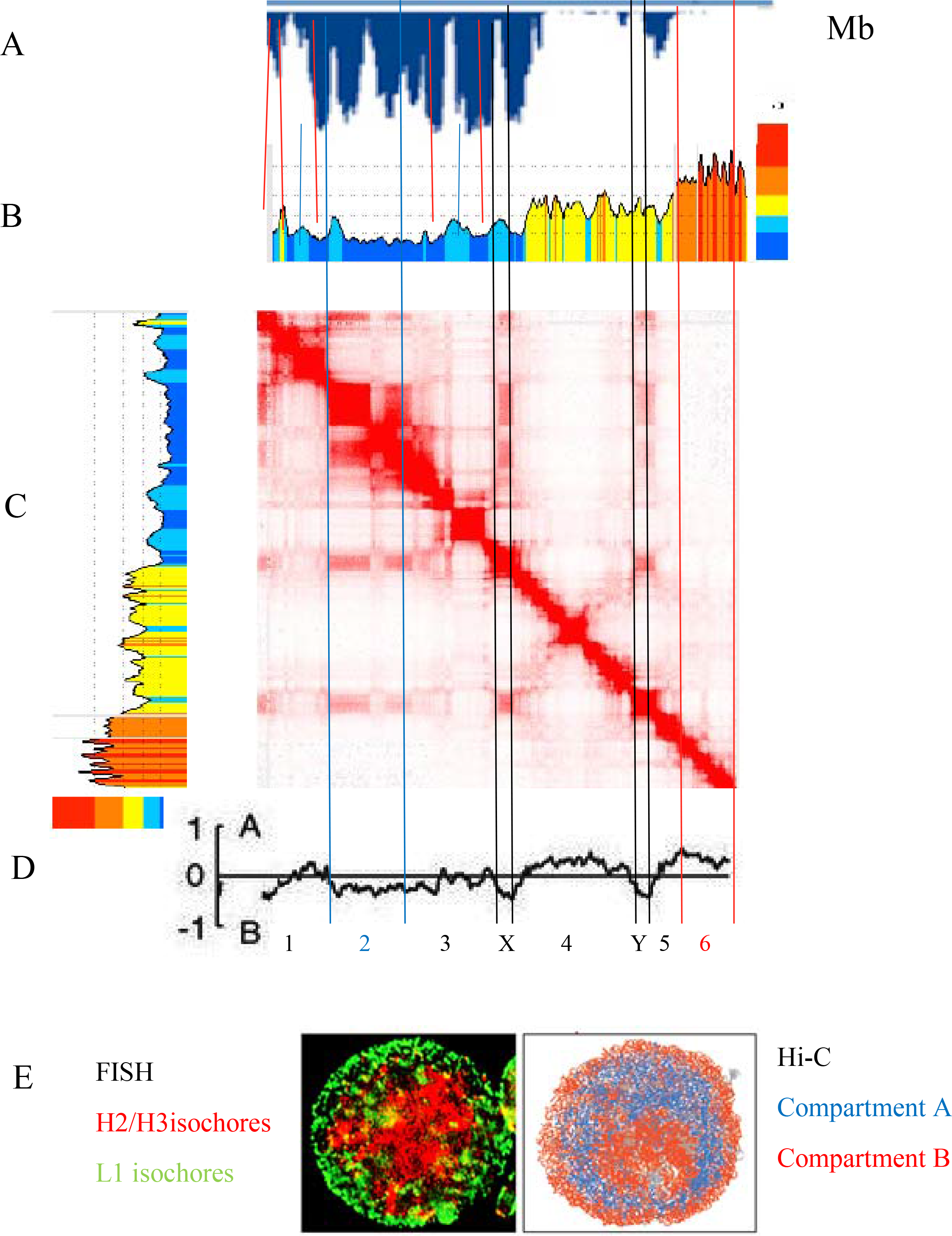
The LAD profile of human chromosome 21, **A** (from ref.40) is compared with 1) the corresponding isochore profile **B**; 2) the heat map of chromatin interactions **C** (from ref. 7 showing several intra-chromosomal interactions of LADs); and 3) the A/B compartment profile **D** (data for mid G1 from ref. 24). For vertical lines, double lines and regions see legend of Fig.1D. in panels A-B red lines link H1 and H2 peaks of inter-LADs. **E.** Results on open and closed chromatin in interphase nuclei as obtained 1) by FISH using H2/H3 and L1 probes on human lymphocytes (red and green signals respectively; yellow signals correspond to the overlap of red and green signals; see ref.22 for details); and 2) by Hi-C on haploid mouse ES cells (25; compartment A blue, compartment B red); in this case, both the nuclear membrane and the nucleolar periphery are in red). Figure 2E was kindly provided by Salvo Saccone.

The results of Fig.2A (as well as similar ones on chromosomes 20 and 19; see Supplementary Fig.S2) show that LADs correspond not only to L1 isochores that represent ~19% of the genome (incidentally, a value not too far from the ~15% involved in “stable contacts”; see ref. 1), but also to L2 isochores (and even to H1 isochores in the rare case of very GC-rich chromosome regions such as those present in chromosome 19).

As far as L2 isochores are concerned, it appears that some of them show a flat profile correspond to LADs and may be called L2^-^ (see, for example, the largest LAD of chromosome 20 in Supplementary Fig.S2B), whereas other ones are in the shape of single peaks (see Figs. 1 A-D and 2A), correspond to interLADs/TADs and may be called L2^+^ (see the following section). The remaining L2 isochores are present as valleys flanked by GC-richer isochores (see Supplementary Fig. S2 and Supplementary Table S1 for the relative amounts of L2 sub-families).

## Isochores and TADs

It should be recalled that the isochores from the five families of the human genome (and other mammalian genomes) are characterized not only by increasing GC levels and different short-sequence frequencies, but also by increasing compositional heterogeneities and increasing frequencies of CpG, CpG islands and Alu sequences (14). Moreover, at the chromatin level, GC increases are correlated with higher bendability (even compared with random sequences having the same composition; 15), higher nuclease accessibility (16), lower nucleosome density (17) and lower supercoiling (18), all properties linked to DNA sequences.

The connection of the isochores of chromosome 21 (as seen in Fig. 1D and in Supplementary Fig. S1), with chromatin loops can be described as follows (see Fig. 2A,B,C): 1) regions 1 and 3 show a series of H1 (a) and L2 (b to f) isochores in which latter case at least some of their single peaks appear to correspond to individual self-interactions; 2) region 2 is the GC-poorest L1 isochore which corresponds to two self-interactions separated by an exceptional GC-poor interLAD; 3) the multi-peak H1 isochores of region 4 correspond to a large interLAD region and to several self-interactions; the two short X and Y sequences, corresponding to LADs, separate region 4 from regions 3 and 5; 4) the small multi-peak H2 isochore (region 5) seems to correspond to a single self-interaction (see below for explanations); 5) a series of H3 GC peaks correspond to a series of self-interactions comprised between the two red lines on the heat map; in this case, the six H3 isochore peaks correspond to at least three chromatin loops (see below).

Two conclusions can be drawn at this point: 1) the two classes of isochores, single-peak and multi-peak, essentially correspond to two classes, single-loop and multi-loop, respectively, of TADs, both of which also show inter-chromosomal interactions (7). It should be mentioned that multi-loop TADs are much more frequent than single-loop TADs as judged from the results of ref.7, and by the fact that H1 and H2 TADs correspond to ~40% of the genome while L2^+^ and H3 TADs only correspond to ~20% of the genome (see Supplementary Table S1); 2) the lack of a one-to-one correspondence between GC peaks and chromatin loops may have two explanations, that are, however, compatible with each other: the first one, the lack of resolution, is demonstrated by the fact that the GC peaks/loops correspondence is improved at high resolution (see Supplementary Figs. S3-S5); the second one is the lack of interaction, *i.e*. the isochore peaks are there, but the loops are not formed (a point discussed later).

## Isochores and chromosome compartments

A higher level of isochore organization was originally discovered (19) by finding that coding sequences are not randomly distributed in the human genome. Later assessments on the increasingly available numbers of genes confirmed a much higher gene density in GC-rich compared to GC-poor isochores. This led to the definition of two “genome spaces” (20,21): the gene-poor, GC-poor “genome desert” (L1 and L2 isochores; the latter being now split into two sub-families, L2^-^ and L2^+^, only the first one belonging to the “genome desert”) and the gene-rich, GC-rich “genome core” (L2^+^, H1, H2, H3 isochores; see Supplementary Table 1 and Supplementary Fig.S6).

In the interphase nucleus of both warm- and cold-blooded vertebrates the chromatin corresponding to the GC-rich, gene-rich isochores was shown to be characterized by an open structure and an internal location, whereas the chromatin corresponding to the GC-poor, gene-poor isochores showed a closed structure and a peripheral location; moreover, large chromosomes showed a polarity because of the preferential locations of GC-rich, gene-rich isochores in telomeric regions, and of GC-poor, gene-poor isochores in centromeric regions (22,23).

Open and closed chromatin domains were later shown to define two different “spatial compartments”, A and B (8), the former one being gene-rich, accessible, actively transcribed, the latter more densely packed. These properties suggest that the A and B compartments may be identified with the open and closed chromatins that correspond to the genome core and to the genome desert respectively. This suggestion is supported 1) by the comparison of sub-compartments (3) with isochore profiles (7) since A1 sub-compartments correspond to H2/H3 isochores (sometimes including flanking isochores from the H1 and even from the L2 family), A2 sub-compartments to H1 and L2 isochores, B1-B3 sub-compartments to L2 and L1 isochores (7); 2) by the comparison of the A and B compartment profile of chromosome 21 (Fig.2D from ref.24) with the corresponding compositional profile (Fig. 2B), the profiles of LADs (Fig. 2A) and the heatmaps of TADs (Fig.2C); indeed, the A compartments correspond to the single L2^+^ peaks of regions 1 and 3 and to the multiple H1, H2, H3 peaks of regions 4 to 6, whereas the B compartments correspond to the L1 isochore of region 2 and to the X and Y sequences; 3) by the similarity of FISH results using H2/H3 and L1 probes (22) with Hi-C results on interphase chromatin of single cells (25).

## Encoding of chromatin domains by isochores

The results from the isochore/chromatin domain connection provide a conclusive evidence for the idea that GC-poor and GC-rich isochores should be visualized as the framework of chromatin architecture or, in other words, as the DNA units that underlie LADs and TADs, respectively (7): 1) by showing in more detail a match between the chromatin domains and the isochores of chromosome 21 (as well as of chromosomes 20 and 19; see Supplementary Fig. S2); 2) by discovering L2^+^ and L2^-^ sub-families on the basis of their correspondence with different chromatin domains (TADs and LADs, respectively); and 3) by generalizing these results to all human chromosomes.

As far as the latter point is concerned, the compositional profiles, the TAD heatmaps and the LAD maps (7) show that: 1) the isochores from the L1 family and the L2^-^ sub-family correspond to LADs in all human chromosomes; 2) L2^+^ peaks emerging from an L1 background and corresponding to interLADs/TADs are also found in other human chromosomes (although less frequently than in chromosome 21); likewise, H3 isochores, that encompass a broad GC range and correspond to interLADs/TADs are present in most human chromosomes; 3) the spikes of the compositional profiles of H1 and H2 isochores of Fig. 1C, that reflect the peaks of Fig. 1D, are regularly present in H1 and H2 isochores from all human chromosomes and correspond to the peaks of the point-by-point profiles (P. Cozzi et al., paper in preparation), as well as to self-interactions of the corresponding chromatin; finally, weak LADs flank centromeres in all chromosomes (see Supplementary Fig.S2). This general match is important in that the only alternative to the encoding proposed here is that the match of thousands of LADs and TADs with the corresponding isochores is just a coincidence (and this cannot be *quia absurdum*).

## Molding of LADs and TADs by isochores

A preliminary remark should be made to stress that the models under discussion here essentially concern the basic evolutionarily stable chromatin domains, since, as it is well known, epigenetic modifications and environmental factors may cause changes in chromatin architecture. Indeed, while the “constitutive” LADs and the TADs discussed here are stable in mammals, chromatin structures and interactions within and between domains may change during differentiation, evolution and even during the cell cycle (26). As already mentioned, this does not prevent the isochore structures typical of chromatin domains still to be present and capable to become the framework of LADs and TADs under the action of appropriate factors.

The present results solve an important open problem, namely the mechanism of formation of chromatin domains. Indeed, LADs should be visualized as chromatin structures corresponding to GC-poor isochores that are flexible, because of the local wider nucleosome spacings linked to the presence of oligo-A sequences; LADs only twist and bend in the three dimensions in order to adapt and attach themselves to (and even embed in) the lamina, which is reassembled after mitosis. This leads to self-interactions (see Fig. 2), as well as to interactions with other LADs from the same chromosomes (7; see, for example, the two X and Y LADs bracketed by black lines in Fig. 2).

In the case of TADs, the GC gradient within each GC-rich isochore peak is accompanied by increasing levels of CpG, CpG islands and Alu sequences that lead to increasing nucleosome spacing, increasing bendability and accessibility, as well as to decreasing supercoiling (12-18). These factors constrain the corresponding chromatin to fold into loops, the tips of the loops corresponding to the highest GC levels, the GC peaks, the architectural proteins (CTCF, cohesin) playing a role in closing the loops and in ensuring loop insulation.

Figs. 3A and 3B display the “isochore molding” models developed here for the formation of LADs and TADs. In these new models, a crucial role is played by isochore sequences, the differential nucleosome binding and the corresponding local (in LADs) or extended (in TADs) flexibilities of the chromatin fibers. While Fig.3B concerns the case of single isochore/single peak/single loop, Supplementary Fig.S5 displays that of single isochore/multiple peaks/multiple loops. Incidentally the “isochore molding” model for TADs is compatible 1) with both the absolute requirement of CTCF binding to close chromatin loops into insulated TADs (27) or the lack of such requirement (28); 2) with an absolute need of cohesin for TAD formation (29); and 3) with the interaction of topoisomerase II beta with cohesin and CTCF at topological domain borders (30).

**Figure 3.**
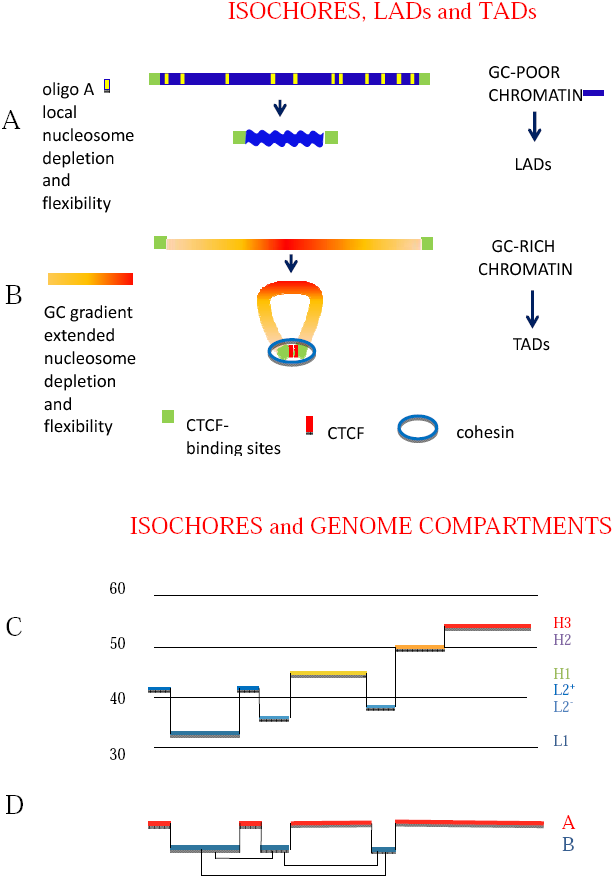
The “isochore molding model” for the formation of LADs and TADs. **A**. A GC-poor chromatin fiber corresponding to an L1 isochore; blue bar) bounded by CTCF binding sites (green boxes) attaches to the lamina, forming a LAD; the wavy profile indicates the three-dimensional physical adaptation by bending and twisting to (and embedding in) the lamina due to local nucleosome depletions associated with oligo-A sequences (yellow bands). **B**. A GC-rich chromatin fiber corresponding to an H3 isochore folds upon itself forming a single-loop TAD bounded by CTCF-binding sites (green boxes); the yellow to red color gradient indicates an increasing GC level with increasing frequencies of CpG islands and Alu sequences. These properties are responsible for the increasing flexibility due to the increasing nucleosome depletion and for the folding. The supercoiling of the loop (see Text) is not shown. Supplementary Fig.S3 displays the case in which a multiple-peak isochore encodes a multiple-loop TAD. **C.** An isochore profile of a chromosome segment in which all families are represented is compared with the corresponding compartment profile to show that while the L1 and L2^-^ isochore correspond to the B compartment, the other isochores correspond to the A compartment. Thin lines and the broken lines connecting B and A compartments, respectively, indicate strong and weak intra-chromosomal interactions.

Going back to the previous models of TAD formation (see Supplementary Fig.S7), the “handcuff” model has two problems: 1) the practical impossibility for the boundary of a given loop to find the other boundary in the very complex nuclear space; and 2) the lack of an explanation for the folding process itself. The main problems of the “extrusion model” are 1) the random location of extrusion start sites that would be, therefore, independent of DNA sequences; and 2) the fact that the source of the energy which is required has not been identified so far. In the “insulation-attraction” model, the problem is that chromatin flexibility is seen as modulated 1) by nucleosome spacing which is not visualized, however, as directly linked to DNA base composition; and/or 2) by other properties.

Now, the “handcuff model” and, even more so, the “insulation-attraction” model can be seen as steps in the direction of the “isochore molding” model and can easily merge into the latter model (compare Supplementary Figs.S7A and S7C with Fig. 3B). In contrast, the problems of the “extrusion model” (see Supplementary Fig.S7B) cannot be solved. Indeed, the predicted random location of extrusion starting sites is contradicted by the fact that the tips of chromatin loops correspond to the highest GC values of isochore peaks. As far the energy source problem, one should consider what happens in the case of the “mitotic memory”, namely in the rapid and precise re-establishment of the original interphase chromatin domains at the exit from mitosis. In the case of the “extrusion model”, the formation of thousands of loops involves the simultaneous attachment of loop extruding factors (the cohesin complex) and extrusion processes that require a source of energy which is still unknown. In the case of the “isochore molding model”, the basic information required for such quick re-establishment is already present in the sequences of isochores. It is obvious that the “extrusion molding” model is incomparably less parsimonious in terms of energy than the “isochore model”.

In conclusion, the “isochore molding” model of chromatin domains represents a paradigm shift compared to the models just discussed since 1) it is essentially based on well-known physico-chemical properties of isochore sequences such as their differential binding of nucleosomes; 2) it provides the same basic explanation for both TADs and LADs; and 3) it leads, in the case of TADs, to a central folding (relative to boundaries) that facilitates the interaction of binding sequences with architectural proteins.

## Encoding of the genome compartments by isochores

Fig.2 shows that in chromosome 21 the A compartments correspond to GC-rich isochores and to single- or multiple-loop TADs, whereas the B compartments correspond to GC-poor isochores and LADs. The alternance of A and B compartments along chromosomes (see Fig.2D, 3C, 3D and ref.31) and the strong intra-chromosomal interactions of LADs at the lamina and the inter chromosomal interactions of TADs in the central part of the nucleus (7) lead to a three-dimensional folding of interphase chromosomes which is also encoded in the sequences of isochores as shown in Fig.3D (apparently this folding is only absent in the maternal chromatin immediately after fertilization; 32). Under these circumstances, it is not surprising that CTCF depletions (27,28) and cohesin removal (29) do not affect the segregation of the active and inactive genome compartments, since they concern a different level of genome organization, that of chromatin domains.

## The genomic code

This definition was originally coined (33,34; see also ref.11) for two sets of compositional correlations 1) those that hold among genome sequences (for instance, between coding and contiguous non-coding sequences and among the three codon positions of genes); and 2) those that link GC-rich and GC-poor isochores with all the structural/functional properties of the genome tested so far (35; see for example Supplementary Table S2 of ref.7). Here, a third crucial point is added, namely the encoding of chromatin domains and of the open/closed compartments by isochores. Strictly speaking, this is the real genomic code, the correlations just mentioned rather reflecting isochore properties.

The genomic code is important in that 1) it leads to an integrated view of the genome; 2) it implies the now well-known existence (i) of “genomic diseases”, namely of diseases due to sequence alteration that do not affect genes or classical regulatory sequences, but sequences that cause regional changes in chromatin structure (36); and (ii) of large-scale genomic differences among individuals due to insertions/deletions, translocations, etc (37); 3) is the strongest, final argument against “junk DNA”.

## Acknowledgements

The author thanks Paolo Ascenzi for hospitality, Fernando Alvarez-Valin (Montevideo, Uruguay), Giacomo Bernardi (Santa Cruz, California, USA), Oliver Clay (Medellin, Colombia), Hector Musto (Montevideo, Uruguay) Salvo Saccone (Catania, Italy) and, especially, Kamel Jabbari (Cologne, Germany) for critical reading, comments and discussions. Caterina Nuvoli and Marta Ritucci provided excellent technical help. This research was supported by the Kimura Prize for Molecular Evolution and Evolutionary Genomics conferred to the author (Tokyo, June 2016).

## References

1. van Steensel B, Belmont A. Lamina-Associated Domains: links with Chromosome Architecture, Heterochromatin, and Gene Repression. Cell, 2017; 169:780–791.

2. Dixon JR, Gorkin DU, Ren B. Chromatin Domains: The Unit of Chromosome Organization. Mol Cell. 2016;62:668–680.

3. Rao SSP, Huntley MH, Durand NC, Stamenova EK, Bochkov ID, Robinson JT, et al. A 3D map of the human genome at kilobase resolution reveals principles of chromatin looping. Cell. 2014;159: 1665–1680.

4. Vietri Rudan M, Hadjur S, Genetic Tailors: CTCF and Cohesin Shape the Genome During Evolution. Trends Genet. 2015; 31:651–660.

5. Fudenberg G, Imakaev M, Lu C, Goloborodko A, Abdennur N, Mirny LA. Formation of Chromosomal Domains by Loop Extrusion. Cell Rep. 2016;15: 2038–2049. http://biorxiv.org/content/early/2015/08/14/024620

6. Sanborn AL, Rao SSP, Huang S-C, Durand NC, Huntley MH, Jewett AI, et al. Chromatin extrusion explains key features of loop and domain formation in wild-type and engineered genomes. Proc Natl Acad Sci USA. 2015;112: 6456–6465.

7. Jabbari K, Bernardi G. An isochore framework underlies chromatin architecture. Plos One 2017 http://dx.doi.org/10.1371/journal.pone.0168023

8. Lieberman-Aiden E, van Berkum NL, Williams L, Imakaev M, Ragoczy T, Telling A, et al. Comprehensive mapping of long-range interactions reveals folding principles of the human genome. Science. 2009;326: 289–293.

9. Macaya G, Thiery JP, Bernardi G. An approach to the organization of eukaryotic genomes at a macromolecular level. J Mol Biol. 1976;108: 237–254.

10. Corneo G, Ginelli E, Soave C, Bernardi G. Isolation and characterization of mouse and guinea pig satellite deoxyribonucleic acids. Biochemistry. 1968. 7: 4373–4379.

11. Bernardi G. Structural and Evolutionary Genomics. Natural Selection in Genome Evolution. Elsevier, Amsterdam. 2004 (reprinted in 2005).

12. Costantini M, Bernardi G. The short-sequence designs of isochores from the human genome. Proc Natl Acad Sci U S A. 2008;105:13971–13976.

13. Tillo D, Hughes TR. G+C content dominates intrinsic nucleosome occupancy. BMC Bioinformatics. 2009, 10:442.

14. Costantini M, Clay O, Auletta F, Bernardi G. An isochore map of human chromosomes. Genome Res. 2006;16:536–541.

15. Vinogradov AE. DNA helix: The importance of being GC-rich. Nucleic Acids Research. 2003. 31:1838–1844.

16. Di Filippo M, Bernardi G. Mapping DNase-I hypersensitive sites on human isochores. Gene. 2008;419:62–65.

17. Fenouil R, Cauchy P, Koch F, Descostes N, Cabeza JZ, Innocenti C, et al. CpG islands and GC content dictate nucleosome depletion in a transcription-independent manner at mammalian promoters. Genome Res. 2012;22: 2399–2408.

18. Naughton C, Avlonitis N, Corless S, Prendergast JG, Mati IK, Eijk PP, et al. Transcription forms and remodels supercoiling domains unfolding large-scale chromatin structures. Nat Struct Mol Biol. 2013;20: 387–395.

19. Bernardi G, Olofsson B, Filipski J, Zerial M, Salinas J, et al. The mosaic genome of warm-blooded vertebrates. Science. 1985, 228:953–958.

20. Bernardi G. The Vertebrate Genome: Isochores and Evolution. Mol Biol Evol. 1993;10: 186–204.

21. Bernardi G. Isochores and the evolutionary genomics of vertebrates. Gene. 2000. 3–17.

22. Saccone S, Federico C, Bernardi G. Localization of the gene-richest and the gene-poorest isochores in the interphase nuclei of mammals and birds. Gene. 2002;300: 169–78.

23. Federico C, Scavo C, Cantarella CD, Motta S, Saccone S, Bernardi G. Gene-rich and gene-poor chromosomal regions have different locations in the interphase nuclei of cold-blooded vertebrates. Chromosoma 2006; 115:123–128.

24. Naumova N, Imakaev M, Fudenberg G, Zhan Y, Lajoie BR, Mirny LA, et al. Organization of the mitotic chromosome. Science. 2013;342: 948–53.

25. Stevens T J, Lando D, Basu S, Atkinson LP, Cao Y, et al. 3D structures of individual mammalian genomes studied by single-cell Hi-C. Nature. 2017; 544:59–64.

26. Nagano T, Lubling Y, Várnai C, Dudley C, Leung W, et al. Cell-cycle dynamics of áchromosomal organization at single-cell resolution. Nature. 2017; 547: 61–67.

27. Nora EP, Goloborodko A, Valton AL, Gibcus J, Uebersohn A, et al. Targeted degradation of CTCF decouples local insulation of chromosome domains from higher-order genomic compartmentalization. BioRxiv 2017. http://biorxiv.org/content/early/2017/01/07/095802

28. Kubo N, Ishii H, Gorkin D, Meitinger F, Xiong X, Fang R, et al. Preservation of Chromatin Organization after Acute Loss of CTCF in Mouse Embryonic Stem Cells. BioRxiv. 2017; http://biorxiv.org/content/early/2017/03/20/118737

29. Schwarzer W, Abdennur N, Goloborodko A, Pekowska A, Fudenberg G, Loe-Mie Y, et al. Two independent modes of chromosome organization are revealed by cohesin removal. bioRxiv. 2016; http://biorxiv.org/content/early/2016/12/21/095802

30. Uusküla-Reimand L, Hou H, Samavarchi-Tehrani P, Rudan MV, Liang M, et al. Topoisomerase II beta interacts with cohesin and CTCF at topological domain borders. Genome Biol. 2016; 17: 1–22.

31. Wang S, Su J-H, Beliveau BJ, Bintu B, Moffitt JR, Wu C -t., et al. Spatial organization of chromatin domains and compartments in single chromosomes. Science, 2016;353: 598–602.

32. Flyamer IM, Gassler J, Imakaev M, Brandão HB, Ulianov SV, et al. Single-nucleus Hi-C reveals unique chromatin reorganization at oocyte-to-zygote transition. Nature. 2017; 544: 110–114.

33. Bernardi G. Le génome des vertébrés: organisation, fonction et évolution. Biofutur. 1990. 94: 43–46.

34. Bernardi G, Bernardi G. Compositional properties of nuclear genes from cold-blooded vertebrates. J. Mol. Evol. 1991. 33:57–67.

35. Bernardi G. Chromosome Architecture and Genome Organization. PLoS One. 2015. 10: e0143739.

36. Bernardi G. The neoselectionist theory of genome evolution. Proc Natl Acad Sci U S A. 2007;104: 8385–90.

37. Costantini M., Bernardi G. Mapping insertions, deletions and SNPs in Venter s chromosomes. Plos One. 2009; 4:1–11.

38. Cozzi P, Milanesi L, Bernardi G. Segmenting the human genome into isochores. Evol Bioinforma. 2015;11: 253–261.

39. Pavícek A, Paces J, Clay O, Bernardi G. A compact view of isochores in the draft human genome sequence. FEBS Lett. 2002;511: 165–169.

40. Kind J, Pagie L, De Vries SS, Nahidiazar L, Dey SS, Bienko M, et al. Genome-wide Maps of Nuclear Lamina Interactions in Single Human Cells. Cell. 2015;163: 134–147.

41. Bernardi G. Misunderstandings about isochores. Part 1. Gene. 2001. 276:3–13.

42. International Human Genome Spacing Consortium, 2001. Initial Sequences and analysis of the human genome. Nature 409, 860–921.

43. Jabbari K, Nürnberg P. A genomic view on epilepsy and autism candidate genes. Genomics. 2015. 108:31–33.

44. Vietri Rudan M, Hadjur S, Genetic Tailors: CTCF and Cohesin Shape the Genome During Evolution. Trends Genet. 2015; 31:651–660.

45. Dolgin E. A loop of faith. Nature. 2017;544: 284–286.

